# Histone deacetylase inhibition by gut microbe-generated short chain fatty acids entrains intestinal epithelial circadian rhythms

**DOI:** 10.1101/2020.06.09.143057

**Authors:** Deborah H. Luzader, Jibraan A. Fawad, Thomas J. Moutinho, Paul G. Mitchell, Kathleen Brown-Steinke, Jesse Y. Zhao, Andrew E. Rosselot, Craig A. McKinney, Christian I. Hong, C. James Chou, Jason A. Papin, Sean R. Moore

**Author notes:** Authors contributed equally (Deborah H. Luzader, Jibraan A. Fawad). **Correspondence:** Sean R. Moore, UVA Child Health Research Center MR-4 Bldg, 409 Lane Rd., Charlottesville, VA 22908, Laboratory: Rooms 2123 and 2127. **AUTHOR CONTRIBUTIONS (using CRediT Taxonomy):** D.H.L.: Conceptualization, Formal analysis, Investigation, Methodology, Project administration, Software, Validation, Writing - original draft J.A.F.: Formal analysis, Investigation, Methodology, Project administration, Validation, Visualization, Writing - original draft T.J.M.: Formal analysis, Methodology, Software, Visualization, Writing - original draft P.G.M.: Investigation, Visualization, Writing - original draft K.B.S.: Investigation, Methodology J.Y.Z.: Software A.E.R.: Investigation, Methodology, Writing - original draft C.A.M.: Investigation C.I.H.: Conceptualization, Funding acquisition, Methodology, Resources, Supervision, Writing - review, & editing C.J.C.: Methodology, Resources J.A.P.: Conceptualization, Funding acquisition, Methodology, Resources, Supervision, Writing - review & editing S.R.M.: Conceptualization, Funding acquisition, Methodology, Project administration, Resources, Supervision, Writing - review & editing.

## Abstract

**Background and aims:** The circadian clock orchestrates ~24-hour oscillations of gastrointestinal (GI) epithelial structure and function that drive diurnal rhythms in the composition, localization, and metabolism of gut microbiota. Here, we use experimental and computational approaches in enteroids to reveal reciprocal effects of microbial metabolites on intestinal epithelial timekeeping by an epigenetic mechanism.

**Methods:** We cultured 3D PER2∷LUCIFERASE and *Bmal1-ELuciferase* jejunal enteroids in media supplemented with sterile supernatants from the altered Schaedler Flora (ASF), a defined murine microbiota. Circadian oscillations of bioluminescent PER2 and *Bmal1* were measured in enteroids cultured in the presence or absence of individual ASF supernatants. Separately, we applied machine learning to ASF metabolic profiles to identify phase-shifting metabolites.

**Results:** Filtrates from 3 of 7 ASF species (ASF360 *Lactobacillus intestinalis*, ASF361 *Ligilactobacillus murinus*, ASF502 *Clostridium* spp.) induced minimal alterations in circadian rhythms, whereas 4 ASF species (ASF356 *Clostridium* spp., ASF492 *Eubacterium plexicaudatum*, ASF500 *Pseudoflavonifactor* spp., ASF519 *Parabacteroides goldsteinii*) induced profound, concentration-dependent phase delays. Random forest classification identified short chain fatty acids (SCFA: butyrate, propionate, acetate, and isovalerate) production as a discriminating feature of “shifters”, i.e., ASF taxa whose metabolites induce phase delay. Experiments with SCFAs confirmed machine learning predictions, with a median phase delay of 6.2 hours. Pharmacological or botanical HDAC inhibitors generated similar phase delays. Further, mithramycin A, an inhibitor of HDAC inhibition, abrogated SCFA-induced phase delays by 20% (*P*<0.05). Key findings were reproducible in human *Bmal1-luciferase* enteroids.

**Conclusions:** Gut microbe-generated SCFAs entrain intestinal epithelial circadian rhythms, in part, by an HDACi-dependent mechanism, with critical implications for understanding microbial and circadian network regulation of intestinal epithelial homeostasis.

## INTRODUCTION

The symbiotic relationship between mammalian hosts and their commensal gut microbes is dynamic and highly dependent on environmental cues^1^. The circadian clock, a key timekeeper in the dance between hosts and their microbes, is a 24-hour biological oscillator that coordinates changes in organismal behavior and physiology in anticipation of environmental fluctuations over the 24-hour day/night cycle. The clock is centrally controlled by the suprachiasmatic nucleus of the hypothalamus, which is entrained by light-dark and sleep-wake cycles^2^. The suprachiasmatic nucleus in turn coordinates peripheral clocks such as the gut regulating tissue-specific, clock-controlled genes^3^. In mammals, the molecular basis of the clock is an autoregulatory transcriptional-translational feedback loop involving CLOCK and BMAL1 heterodimers. CLOCK and BMAL1 are transcription factors that are highly expressed during the day and control the expression of core clock genes, *Period* (*Per1*, *Per2*, *Per3*) and *Cryptochrome* (*Cry1*, *Cry2*), which are negative elements that are expressed during the dark phase^4^. As PER and CRY proteins accumulate, they form complexes by directly binding to CLOCK/BMAL1 subsequently inhibiting their transcription^5–7^. PERIOD2 (PER2) has been implicated as an important negative element in feedback loops that generate circadian rhythms^5,8,9^.

Numerous intestinal epithelial functions are regulated by the circadian clock, including nutrient absorption, enterocyte proliferation, and metabolism^10^. Day-night cycles have played an important evolutionary role in coordinating when food is consumed and digested^11^. Unsurprisingly, circadian disruption is associated with several gastrointestinal disorders such as gastric ulcers^12^, diarrhea^13^, obesity^14^ and flares of inflammatory bowel disease^15^. Recent work has uncovered an intimate connection between the host mammalian circadian clock and diurnal oscillations of composition, metabolic functions, and localization of the intestinal microbiome, which plays a role in the entrainment and modulation of host circadian signaling^16^. These microbial rhythms in turn influence diurnal host gene expression in the intestine and the liver^17^. Rhythmic expression of host toll-like receptors (TLRs) also entrain microbiota signaling in arrhythmic microbiota, further demonstrating this bidirectional relationship^18^. Diet plays an important role in the maintenance of circadian rhythms as well. Consumption of a high-fat diet in animal models has been shown to alter the period of the central clock indicating diet influences circadian rhythm signaling^19^.

Several studies have focused on circadian rhythms of the liver and its crosstalk between the gut microbiome^17,20,21^. More recent work has identified a pivotal role for intestinal microbiota in programming diurnal metabolic rhythms in intestinal epithelial cells through specific histone deacetylases (HDAC) using *in vivo* approaches^22^. Previously, we have shown that intestinal epithelial organoids (enteroids) provide an ideal experimental platform for elucidating circadian regulation of intestinal epithelial homeostasis^23,24^. Here, we combine this approach with systems metabolomics to demonstrate: 1) entrainment of the host circadian clock by metabolites from the altered Schaedler Flora (ASF), a simplified mouse gut microbiome, 2) machine learning identification of short chain fatty acid (SCFA) production as a key discriminating feature of “shifters” (ASF taxa whose metabolites entrain circadian rhythms), and 3) an HDAC inhibitory (HDACi) dependent mechanism by which SCFAs induce circadian phase delay.

## MATERIALS AND METHODS

### Altered Schaedler Flora Strain Information

All strains used in this study originate from the altered Schaedler Flora, a defined mouse gut microbiota. ASF strains were a gift to the Papin lab from Drs. Michael Wannemuehler and Gregory Phillips. The members used in this study are: *Clostridium* sp. ASF356, *Lactobacillus intestinalis* ASF360, *Ligilactobacillus murinus* ASF361, *Eubacterium plexicaudatum* ASF492, *Pseudoflavonifractor* sp. ASF500, *Clostridium* sp. ASF502, and *Parabacteroides goldsteinii* ASF519. *Mucispirillum schaedleri* ASF457 was excluded from the study due to lack of detectable growth in the experimental medium. Stock vials for all ASF strains were maintained at −80°C in 50% glycerol. All strains were grown in an anaerobic chamber (Shellab BACTRONEZ, Sheldon Manufacturing, Inc., Cornelius, Oregon, USA) filled with mixed anaerobic gas (5% CO_2_, 5% H_2_, 90% N_2_). Anaerobic conditions were ensured using palladium catalysts (baked at 75°C outside the chamber and rotated daily when first entering the chamber) and anaerobic indicator strips (Oxoid, Basingstoke, UK).

### Media Formulations

Full media formulations can be found in supplemental methods. Supplemented Brain–Heart Infusion medium (BHI) and modified Lennox Broth medium (mLB) were used to grow ASF bacterial species. Murine enteroids were grown in murine enteroid growth medium (mEGM)prepared using 70% L-WRN conditioned medium mixed with 30 % basic minigut medium. Human *Bmal1-luciferase (Bmal1-luc)* enteroids were grown in Stemcell IntestiCult™ Organoid Growth Medium (Human) (OGM) (CAT#06010).

### Bacterial Species Growth and Supernatant Collection

ASF species were grown as previously described^25^. Agar plates containing supplemented BHI media were equilibrated inside an anaerobic chamber for a minimum of 24 hours. Strains were inoculated from frozen stock to grow until dense on these agar plates. Plates were incubated for 3-9 days at 37°C before being used to start cultures in 50 mL mLB broth in 500mL glass flasks covered with a Breathe-Easy membrane (Diversified Biotech, Dedham, Massachusetts, USA). After incubating for 72 hours at 37°C, these cultures were transferred to 50 mL conical tubes, sealed and transferred out of the chamber to be centrifuged at 15,000 xg for 5 minutes. After centrifugation, the supernatant was poured off and filter-sterilized (0.22 um pore size, mixed cellulose ester filter) to acquire sterile bacterial supernatant.

### Isolation of Murine PER2∷LUCIFERASE and *Bmal1-ELuc* enteroids

PER2∷LUCIFERASE (PER2∷LUC) mice^26^ on a C57Bl/6J background were obtained from Jackson Laboratories. *Bmal1-ELuc* mice were received from David Welsh’s lab at UCSD given as gifts from Yoshihiro Nakajima’s lab at National Institute of Advanced Industrial Science and Technology^27^. Mice were housed in the institution’s vivarium facility with an ambient temperature of 22°C and a 14-hour light: 10-hour dark cycle. Mice were euthanized using inhaled isoflurane followed by cervical dislocation according to the guidelines of the University of Virginia’s Institutional Animal Care & Use Committee.

Enteroids were isolated as previously described^28^. Jejunal tissue was dissected from mice, and crypts were released by incubating tissue in PBS containing 2mM EDTA at 4°C for 30 minutes. Isolated crypts were counted and pelleted. A total of 100 crypts were mixed with 40µl of Matrigel™ (BD Biosciences, San Jose, CA, USA) supplemented with recombinant growth factors 50ng/mL Noggin (Peprotech, Rocky Hill, NJ, USA) and R-Spondin (Peprotech, Rocky Hill, NJ, USA), and 25ng/ml of murine EGF. Enteroids were plated in a 24-well tissue culture plate (Costar® 24-well Clear TC-treated Multiple Well Plates) and were grown in culture for 7 days before being passaged into additional wells or plated in 35mm dishes for bioluminescence experiments.

### Human *Bmal1-luc* enteroid generation

Human intestinal enteroids (HIEs) were transduced with pABpuro-BluF (Addgene) plasmid DNA containing 1kb of the mouse *Bmal1* promoter fused to luciferase (*Bmal1-luciferase*)^29^. Details of the HIE line and transduction protocol are provided in the supplemental methods.

### Bioluminescence Recording

Murine PER2∷LUC and *Bmal1*-E*Luc* enteroids or human *Bmal1-luc* enteroids were passaged and plated on 35 mm dishes supplemented with 2mL of either mEGM or IntestiCult OGM respectively. The next day, PER2∷LUC, *Bmal1*-E*Lluc* or *Bmal1-luc* expression was synchronized by adding 50% FBS for 2 hours^32^. After synchronization, medium was replaced with appropriate enteroid medium, supplemented with either ASF filter-sterilized supernatants or microbial metabolites sodium acetate, sodium formate, sodium butyrate, isovaleric acid, sodium propionate, choline chloride, nicotinamide, or L-glutamate (GlutaMAX, Gibco). PER2∷LUC, *Bmal1*-E*Lluc* or *Bmal1*-*luc* abundance was measured in real-time by bioluminescence recording in a Kronos Dio™ AB-2550 incubating luminometer (ATTO Corporation) for 72 hours.

### Data Analysis and Processing

R code was created to analyze, filter out noise, and normalize detrended data obtained from the Kronos Dio™ AB-2550 incubating luminometer (ATTO Corporation). Using peak, zero, and trough identification, the instantaneous period and amplitude of the circadian rhythms were accurately calculated and interpolated in this study. Additionally, phase shifts were calculated when comparing two rhythms (i.e. control vs. experimental). Code is available at: https://github.com/Tjmoutinho/GI_circadian_rhythms. Differences in amplitude and period between either ASF supernatant or metabolites were compared with 25% mLB or media vehicle controls, respectively, by a Mann-Whitney U test.

### Machine Learning with Metabolomics Data

Metabolomics of ASF species were used to identify metabolites associated with circadian shifts^25^. The R package AUCRF (v. 1.1) was used to generate a Random Forest (RF) model to classify samples according to whether they were shifters or non-shifters. The normalized metabolomics data were used to train the model by treating all metabolites with a median greater than zero and greater than the non-shifter median as features. Features meeting the described requirements were produced in the majority of samples and therefore allowed for *in vitro* investigation, whereas all other features were removed from the analysis. During the creation of the optimal RF, 5-fold cross validation was used. The optimal forest had 2000 trees with a node size of 5, and classes were manually weighted (using the ‘classwt’ parameter) to account for the imbalance between the shifter and non-shifter classes.

## RESULTS

### Metabolites from ASF species differentially modulate the intestinal epithelial clock

Incubating PER2∷LUC enteroids with bacterial supernatants from ASF356, ASF492, ASF500 and ASF519 significantly altered circadian phase (**Figure 1B**) while supernatants from ASF360, ASF361 and ASF502 did not (**Figure 1A**). Mann-Whitney U test indicated significant phase delays at both 10% and 25% concentrations of ASF356 (10%, median=7.09 hours, *P*<0.05; 25%, median=10.89 hours, *P*<0.05), ASF500 (10%, median=2.2 hours, *P*<0.05; 25%, median=7.67 hours, *P*<0.05), and ASF519 (10%, median=8.62 hours, *P*<0.05; 25%, median=14.3 hours, *P*<0.05) relative to enteroids incubated with 25% mLB alone (**Figure 2**). Additionally, a 25% concentration of ASF492 induced a significant phase delay (median=3.37 hours, *P*<0.05) (**Figure 2**). We observed similar findings in murine *Bmal1*-E*Luc* enteroids, with significant phase delays at 10% concentrations of ASF356 (median=10.51 hours, *P*<0.05) relative to enteroids exposed to 25% mLB alone (**Supplementary Figure 1**).

**Figure 1.**
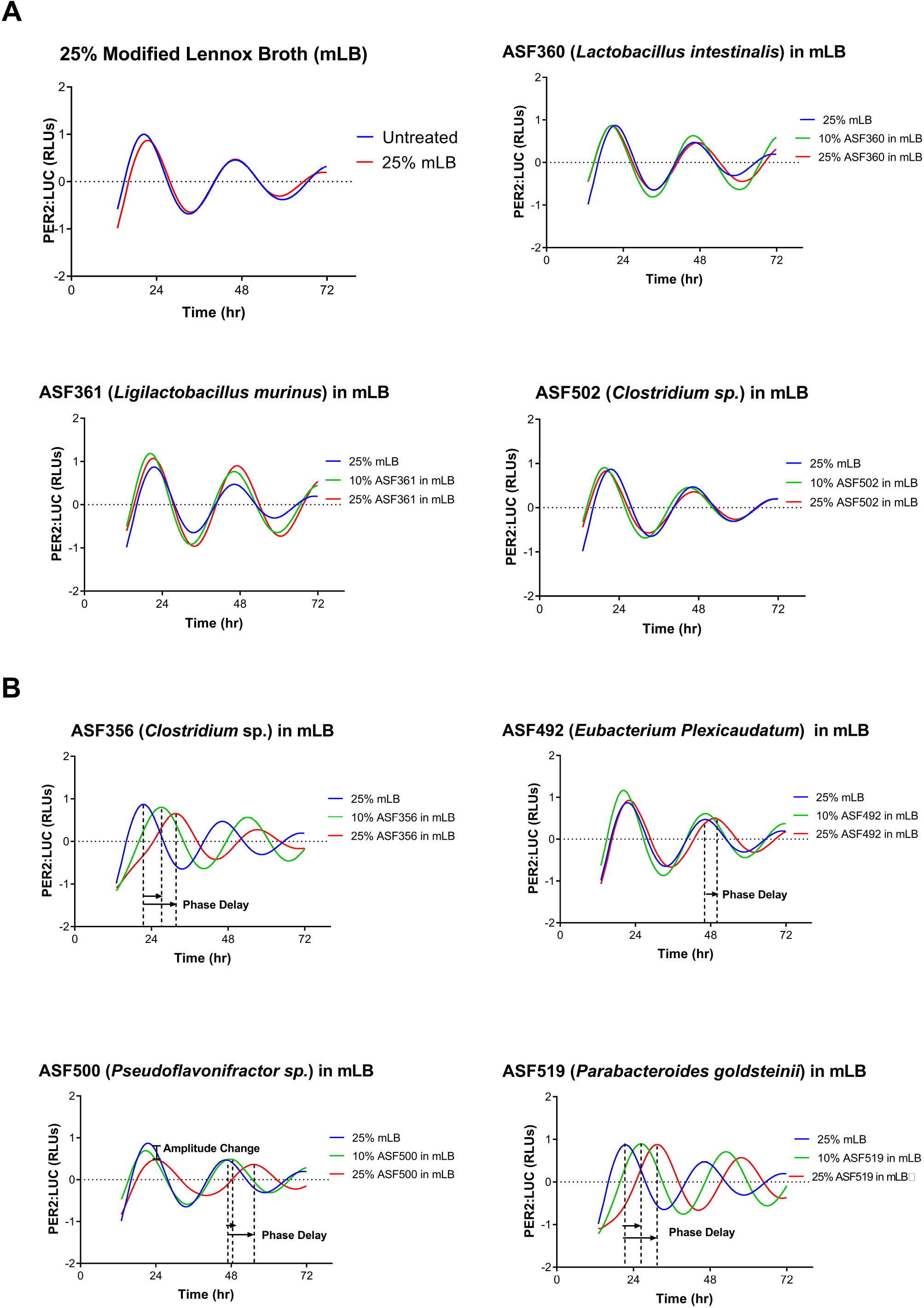
Representative PER2∷LUC enteroids displaying PER2∷LUC oscillations over 72 hours. (A) 25%mLB and ‘non-shifter’ group of ASF species (ASF360, ASF361, ASF502) do not show alterations in PER2∷LUC oscillations. (B) ASF ‘shifter’ group (ASF356, ASF492, ASF500, ASF519) causes phase delays in PER2∷LUC oscillations represented as a forward shift in the PER2∷LUC waveform (*P*<0.05 by Mann-Whitney *U* test, n=3 samples per taxa). ASF500 results in decreased amplitude of PER2∷LUC shown as a decrease in height of PER2∷LUC waveform.

**Figure 2.**
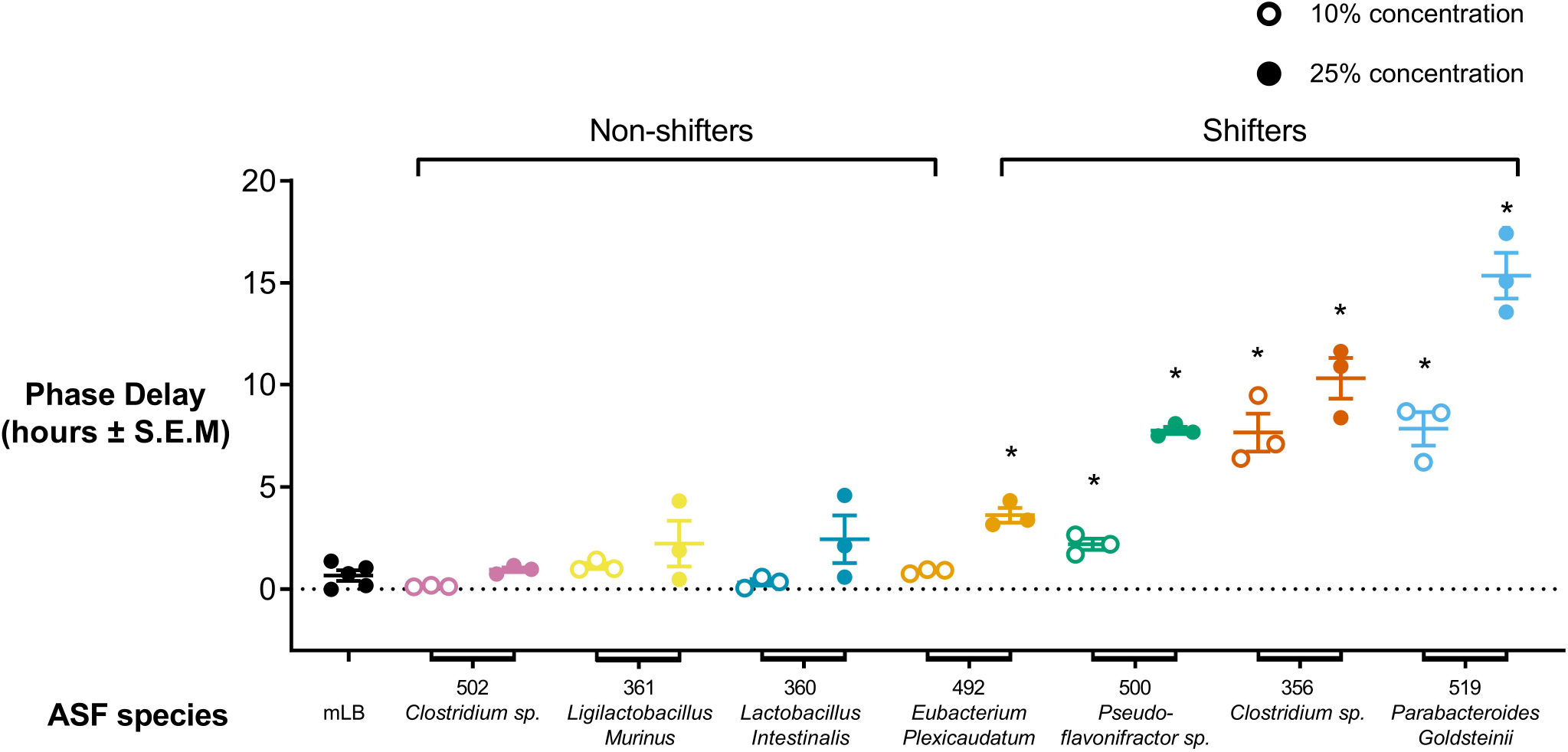
Median phase delay (represented by middle horizontal line of samples) due to mLB or ASF bacterial supernatant calculated as a forward shift in PER2∷LUC oscillation compared to an untreated control. ASF ‘shifters’ classified as ASF taxa causing statistically significant phase delays of PER2∷LUC oscillation as compared to a 25% mLB control (\**P*<0.05, Mann Whitney *U* test).

At a 25% concentration, ASF500 dampened the average amplitude of PER2∷LUC abundance (median= 0.395, *P*<0.05) when compared with 25% mLB alone (median=0.7032) (**Supplementary Figure 2A**). Additionally, the 25% concentration of ASF500 caused significant period changes in PER2∷LUC abundance (median= 27.96 hours, *P*<0.05) when compared with 25% mLB alone (median=25.52 hours, p>0.05) (**Supplementary Figure 2B**). Among murine *Bmal1-ELuc* enteroids no significant modulations in period or amplitude were observed (**Supplementary Figure 3**). With these findings we conclude that phase delays caused by ASF metabolites are primarily a result of PER2 or *Bmal1* abundance alteration and not due to changes in circadian period.

We hypothesized specific microbe-generated metabolites present in ASF supernatants play an active role in entrainment of host circadian rhythms. To test this hypothesis, we analyzed metabolomics data from ASF bacterial supernatants^25^ to identify specific metabolites as key determinants of circadian entrainment.

### Machine learning predicts metabolites contributing to circadian phase delay

Using machine learning applied to a set of 53 detectable metabolites, we identified the top metabolites most likely to contribute to a shift in circadian rhythm. We identified eleven metabolites as candidates (**Figure 3A**). By identifying top candidates we were able to narrow our experimental search for metabolites correlated with circadian phase delays. Using a feature-reduced random forest model we identified three microbial metabolites (acetate, butyrate and isovalerate) that were most highly associated with the circadian shift (**Figure 3B&C**). These three metabolites alone gave an accuracy of 98% in predicting which group each ASF belongs to (shifter vs. non-shifter). Our model was not overfit to the dataset according to the cross validation results: both the cross validation accuracy (cvAUC) and optimal model accuracy (OOB-AUCopt) are nearly the same.

**Figure 3.**
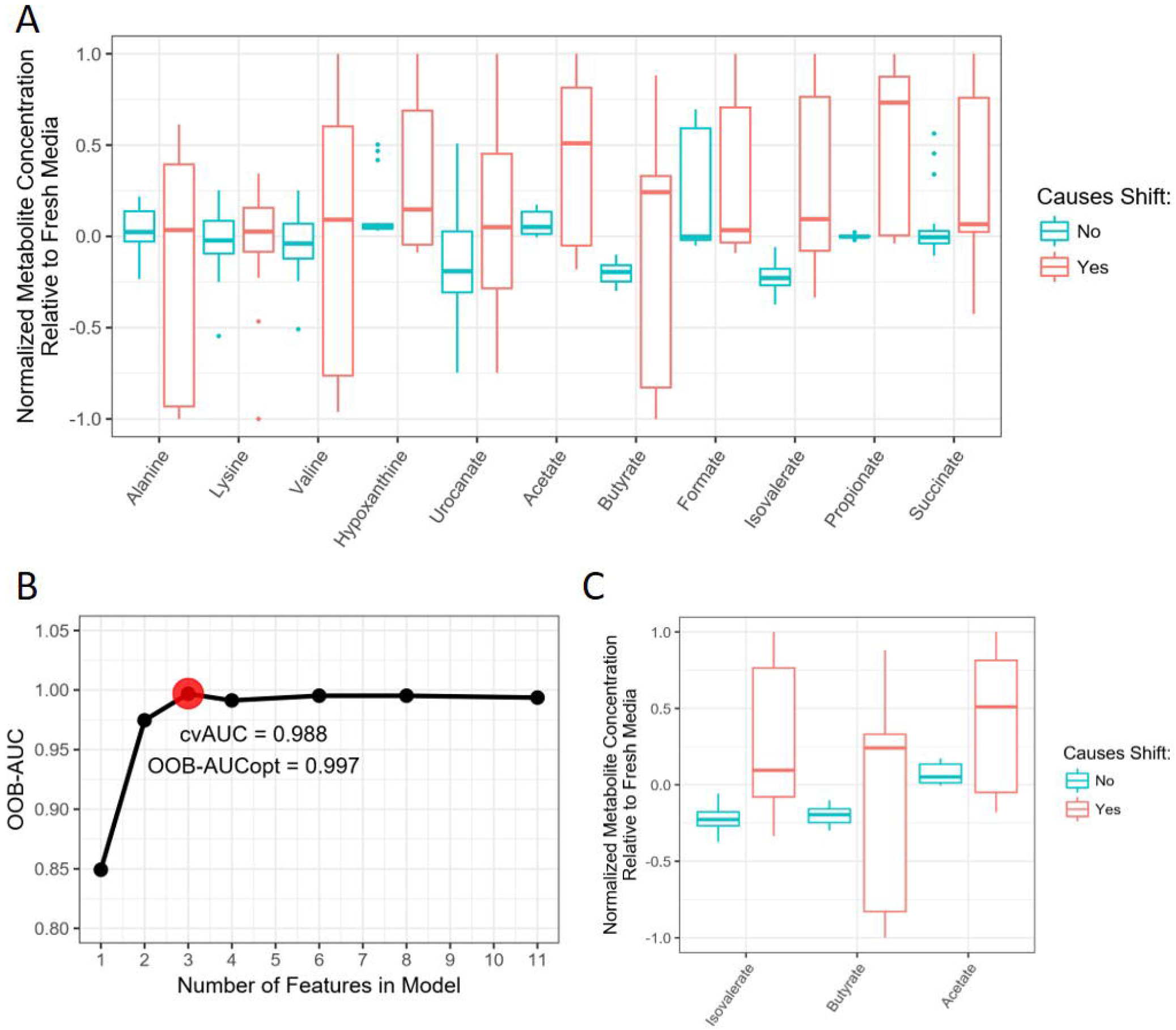
A) Eleven out of 53 detectable metabolites have a greater concentration in shifters (356, 492, 500, 519) compared to non-shifters (360, 361, 502). Additionally, the metabolites included in this group required having a median value greater than zero. These represent the metabolites that are experimentally testable to assess if their presence influences the circadian rhythm of enteroids in culture. B) With a 5-fold cross-validation, using 20 repetitions, the reported accuracy of using random forests to discriminate between the shifter and non-shifter group is 98% with the use of only three features. C) The three essential features for discriminating the groups are displayed in the boxplots. The value plotted on the y-axis is the normalized metabolite concentration relative to fresh media, therefore values above zero indicate that the metabolite was produced by the bacteria in culture. The middle lines in the boxplots represent the median value.

### Host microbe-generated short-chain fatty acids alter host circadian clock in the GI tract

Using the machine learning methods above, we shortlisted specific bacterial metabolites within ASF supernatants as likely contributors to alterations in the circadian clock. SCFAs arose as a key discriminating feature of phase shifting ASF species. We then tested the SCFAs acetate, butyrate, propionate and isovalerate as well as other bacterial metabolites such as glutamine, choline and nicotinamide to validate model predictions.

In PER2∷LUC enteroids 10mM acetate (median=3.72 hours, *P*<0.05), 0.5mM and 1mM isovaleric acid (0.5mM, median=5.1 hours, *P*<0.05; 1mM, median=8.1 hours, *P*<0.05), 1mM and 10mM propionate (1mM, median=3.26 hours, *P*<0.05; 10mM, median=11.74 hours, *P*<0.05), and 1mM butyrate (median=13.35 hours, *P*<0.05) caused significant phase delays in PER2∷LUC abundance when compared with untreated controls (**Figure 4**, **5**). Formate, glutamine, choline and nicotinamide did not modulate PER2∷LUC abundance (**Supplementary Figure 4**). Additionally, enteroids treated with 10mM hydrochloric acid (HCl) (median= −2.25 hours, *P*<0.05) and 10mM sodium hydroxide (NaOH) (median= −22.02 hours, *P*<0.05) contributed to phase advances of PER2∷LUC abundance when compared with an untreated control (**Figure 5**, **Supp Figure 4**). This signifies that pH changes are not responsible for phase delays due to these microbial metabolites.

**Figure 4.**
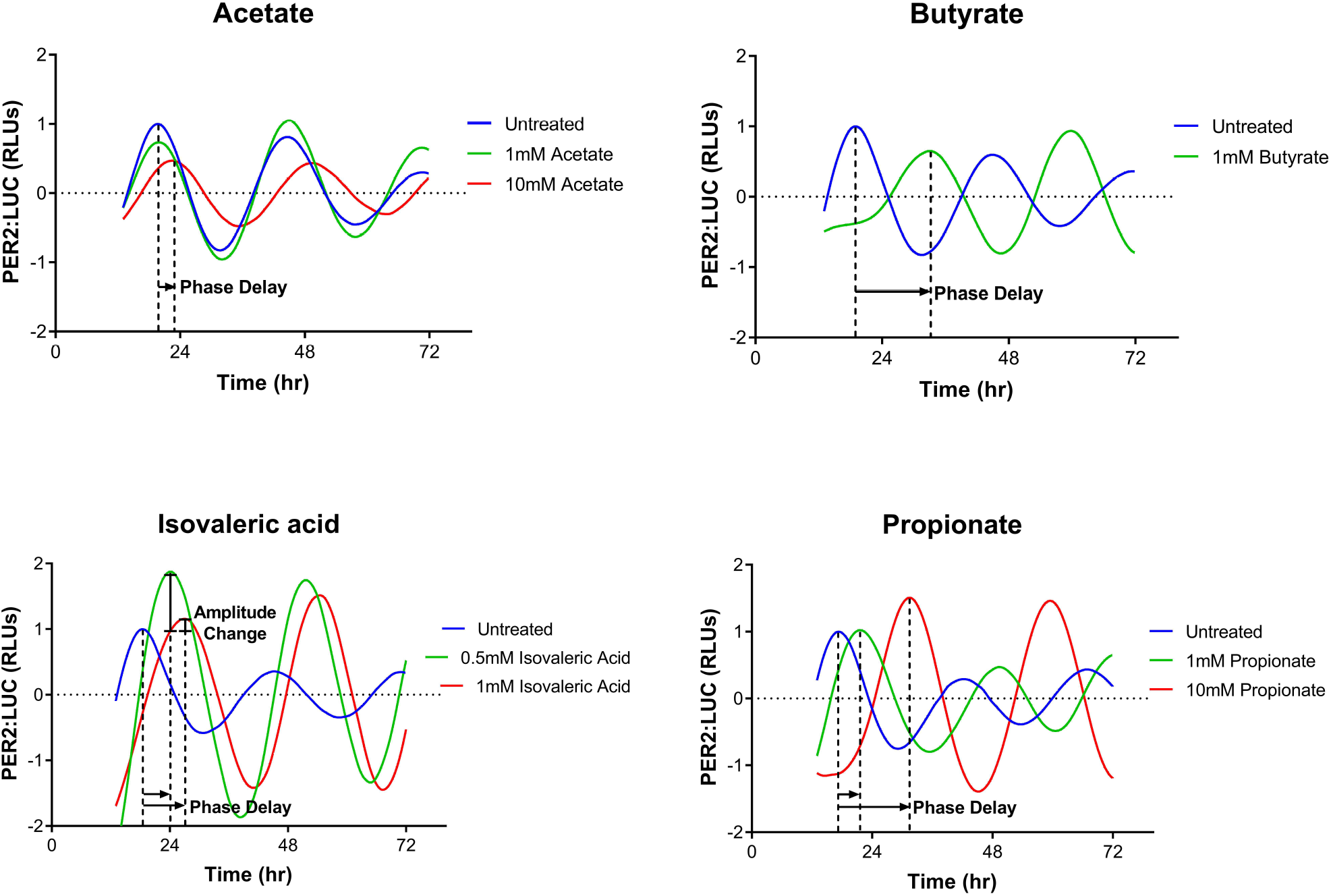
Representative PER2∷LUC enteroids displaying PER2∷LUC oscillations over 72 hours. Acetate, butyrate, isovaleric acid and propionate cause phase delays in PER2∷LUC oscillations represented as a forward shift in the PER2∷LUC∷LUC waveform (*P*<0.05 by Mann-Whitney *U* test, n=3 samples per metabolite). Isovaleric acid results in increased amplitude of PER2∷LUC shown as an increase in height of PER2∷LUC waveform.

**Figure 5.**
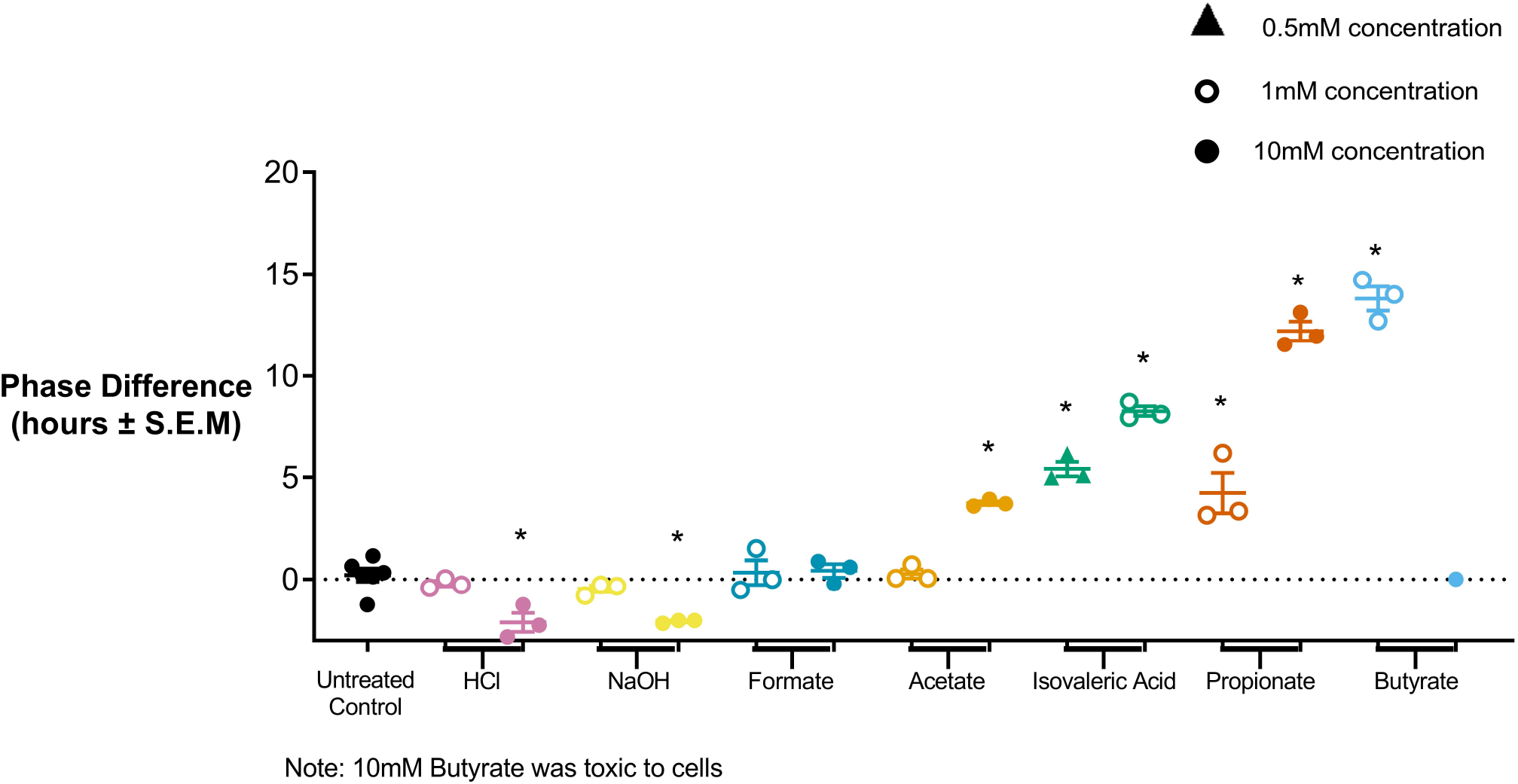
Median phase delay (represented by middle horizontal line of samples) due to ASF-generated metabolites calculated as a forward shift in PER2∷LUC oscillation compared to an untreated control. Shifting metabolites are those causing statistically significant phase delays (forward shift) or phase advances (backward shifts) in PER2∷LUC oscillation compared to untreated controls (\**P*<0.05, Mann Whitney *U* test). 10mM Butyrate was toxic to cells, does not represent a phase shift.

Similar to PER2∷LUC enteroids, *Bmal1-ELuc* enteroids exposed to 10mM acetate (median=3.85 hours, *P*<0.05), 1mM propionate (median=7.61 hours, *P*<0.05), 10mM propionate (median=15.87 hours, *P*<0.05), and 1mM butyrate (median=11.08 hours, *P*<0.05) exhibited significant phase delays in *Bmal1-ELuc* expression relative to untreated controls (median=0.45 hours) (**Supplementary Figure 5**).

Interestingly, 0.5mM isovaleric acid (median=1.069, *P*<0.05) and 1mM isovaleric acid (median=1.39, *P*<0.05) generated an increase in the average amplitude of PER2∷LUC abundance relative to untreated controls (median=0.732), whereas 10mM NaOH (median=0.45, *P*<0.05) and 1mM formate (median=0.620, *P*<0.05) produced a decrease in average amplitude (**Supp Figure 6A**). *Bmal1-Eluc* enteroids exposed to 10mM formate (median=0.78, *P*<0.05), 10mM acetate (median=0.507, *P*<0.05), 1mM propionate (median=0.291, *P*<0.05), and 1mM butyrate (median=0.407, *P*<0.05) displayed significant amplitude modulation of *Bmal1-ELuc* expression versus untreated controls (median=0.66, p>0.05) (**Supp Figure 7A**).

10mM acetate(median=26.35, *P*<0.05), 10mM propionate (median=27.13 hours, *P*<0.05) and 1mM butyrate (median=26.59, *P*<0.05) caused significant increases in the period of PER2:LUC abundance, whereas 10mM HCl (median=23.7 hours, *P*<0.05) and 10mM NaOH (median=24.2 hours, *P*<0.05) caused significant decreases relative to untreated controls (median=25.7 hours) (**Supplementary Figure 6B**). Only 1mM propionate (median period=22.78 hours, *P*<0.05) elicited a significant period shift in enteroid *Bmal1-ELuc* expression when compared with untreated controls (median period=24.27 hours, *P*<0.05) (**Supp Figure 7B**).

### Histone deacetylase inhibition causes phase delays in host circadian clock

An important downstream effect of circadian rhythms in generating rhythmic gene expression is via histone modifications to chromatin^20,33–35^. Recent studies have demonstrated the role of the microbiota in programming diurnal rhythms in the small intestine through histone deacetylases^22^. Interestingly, SCFAs such as butyrate, propionate and isovalerate have potent HDACi activity, suggesting that HDAC inhibition may be an important mechanism of host circadian rhythm entrainment by the microbiota.

To determine whether inhibition of histone deacetylases may play a role in altering host circadian rhythms, we first exposed enteroids to the known HDACis trichostatin (TSA)^36^, SAHA^36^, LP-411^41^, MS275^43^, and HDAC1-2^42^ inhibitor. Additionally we tested sulforaphane^37,38^ (SFN), an isothiocyonate derived from glucosinolates in the *Brassicaceae* family (e.g. cabbage, radishes). SFN is a diet-derived, microbially activated compound with putative HDACi activity^39^.

Co-incubating PER2∷LUC enteroids with 5uM TSA and 1uM SAHA produced a pattern of PER2∷LUC abundance strongly resembling phase delay patterns seen with 1mM of butyrate over the initial 10 hours (**Figure 6A**). This finding suggests that the HDACi activity of butyrate plays a role in circadian rhythm entrainment. However, TSA and SAHA proved to be toxic to PER2∷LUC enteroids beyond 15 hours of exposure, therefore leading to a search for less potent, yet bioactive HDACis such as LP-411^40^, HDAC1-2 inhibitor^41^, and MS275^42^.

**Figure 6.**
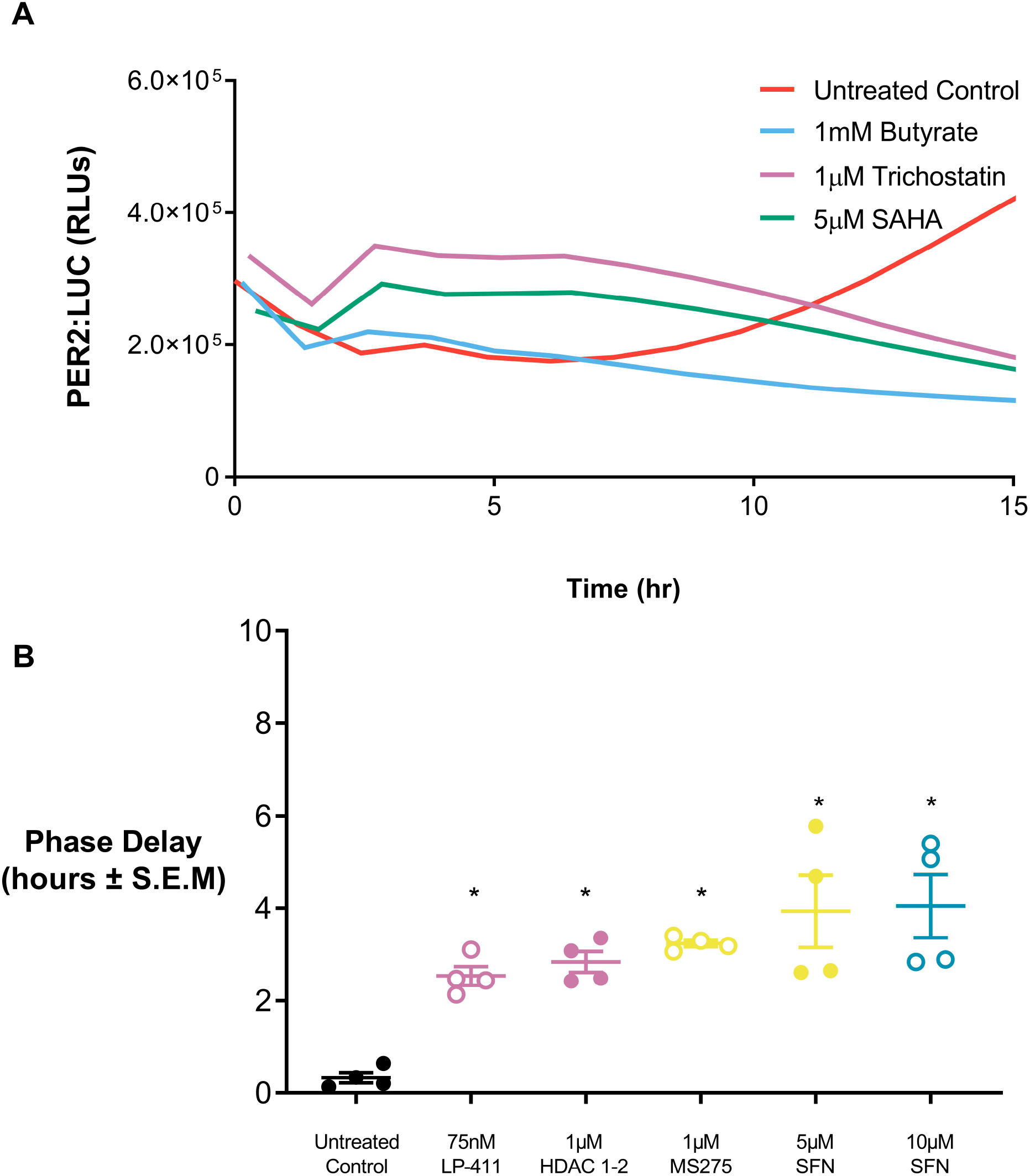
HDAC inhibitors cause significant phase delays in PER2∷LUC abundance. (A) Representative figure showing HDAC inhibitors TSA and SAHA resulting in phase delay patterns of PER2∷LUC similar to phase delay due 1mM butyrate. (B) HDAC inhibitors LP-411, HDAC1-2, MS275 and sulforaphane (SFN) cause significant phase delays in PER2∷LUC abundance compared to an untreated control (\**P*<0.05, Mann Whitney *U* test).

Enteroids treated with 75nM LP-411 (median=2.454 hours, *P*<0.05), 1uM MS275 (median=3.242 hours, *P*<0.05), HDAC1-2 inhibitor (median=2.783 hours, *P*<0.05) and 5uM and 10uM concentration of SFN (5uM, median=3.674, *P*<0.05; 10uM, median=3.979; *P*<0.05) all showed significant phase delays when compared to untreated controls (**Figure 6B**). Taken together, our findings strongly suggest HDACi by microbially produced metabolites is a key mechanism by which gut microbiota entrain intestinal epithelial circadian rhythms.

### SCFA and microbial metabolites alter the host clock through histone deacetylase (HDAC) inhibition

To confirm the role that HDAC inhibition plays in the time-keeping mechanism between the host and the microbiota we used mithramycin A, an inhibitor of Sp1/Sp3 transcription^43^. Inhibition of Sp1/Sp3 transcription abrogates HDAC inhibitor induction of immediate early genes such as PER1 and PER2^32^. Exposure of enteroids to 1mM butyrate supplemented with 0.1 uM of mithramycin A (median phase delay=11.55 hours, *P*<0.05) demonstrated significantly reduced phase delay in PER2 expression when compared to enteroids exposed to 1mM of butyrate alone (median phase delay=13.35 hours) (**Figure 7**). This finding supports our hypothesis that HDAC inhibition is a key intermediary in microbe-derived SCFA induced phase delays.

**Figure 7.**
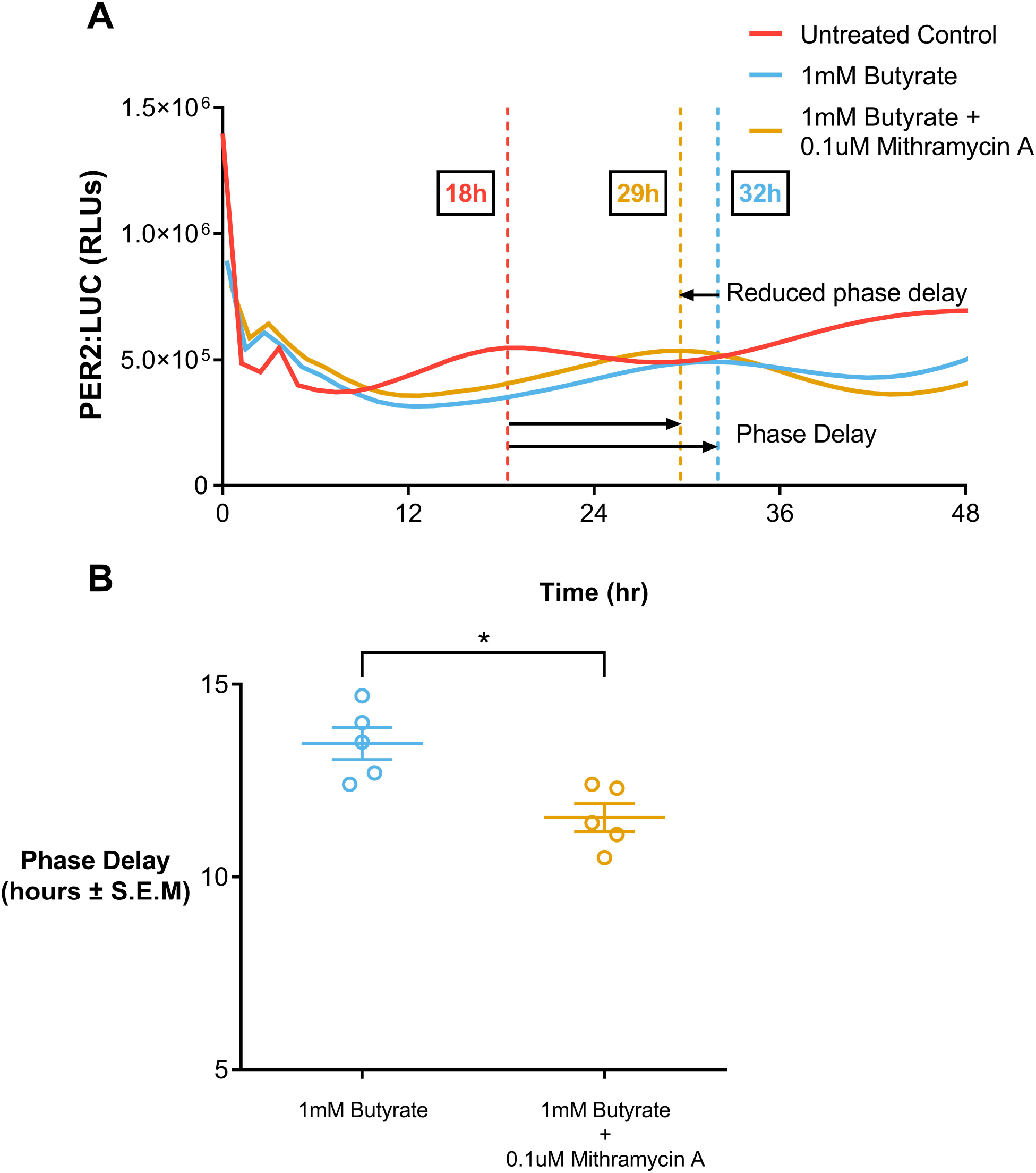
Mithramycin A, an inhibitor of HDAC inhibition, abrogates butyrate induced phase delay of PER2∷LUC abundance. (A) Representative oscillations showing a 3-hour reduction in butyrate induced phase delay of PER2∷LUC abundance in the presence of mithramycin A. (B) Reduction in butyrate-induced median phase delay (represented by middle horizontal line of samples) of PER2∷LUC oscillations due to mithramycin A (\**P*<0.05, Mann Whitney *U* test).

### Human *Bmal1-luc* enteroids display modest phase delays when exposed to microbial metabolites

To explore the reproducibility of our findings from murine enteroids in non-transformed human intestinal epithelial cells, we generated human *Bmal1-luc* enteroids. After confirming *Bmal1* rhythmicity, we then investigated the effects of a potent ASF shifter, ASF356, and propionate on *Bmal1* oscillations. Exposure to 25% ASF356 media (median=1.3 hours, P>0.05) and 10mM propionate (median=2.2 hours, P>0.05) (**Supplementary Figure 8**) produced modest phase delays of *Bmal1* abundance that were not statically significant.

## DISCUSSION

In these *ex vivo* intestinal organoid studies of microbial metabolites and intestinal epithelial circadian timekeeping, we used a combination of experimental methods and systems metabolomics to demonstrate: 1) entrainment of the host circadian clock by metabolites from specific taxa of the altered Schaedler Flora (ASF), 2) machine learning identification of short chain fatty acid (SCFA) production as a key discriminating feature of “ASF shifters”, and 3) an HDACi mechanism by which physiological levels of SCFAs induce circadian phase delay in enteroids. Previous studies have identified circadian oscillations in the liver metabolome,^21,44^, gut microbiota and microbial metabolites^16,17^, and *in vivo* entrainment of the gut circadian clock by short chain fatty acid HDAC inhibition.^22^ Our study extends these important findings by establishing and exploiting a real-time organotypic model of host intestinal circadian rhythms in relation to a complete but simplified microbiome.^23,24^

To study complex interactions between the intestinal epithelium and the microbiome in relation to the clock, we leveraged a well-curated metabolomic dataset from a simplified and defined mouse microbiome, the altered Schaedler Flora (ASF)^25^. ASF species prove a uniquely useful model for understanding host-microbe interactions. Mice colonized with this community can remain stably colonized for multiple generations while promoting normal organ physiology and immune system function^45^. Additionally, the members of the ASF can be cultured *in vitro*, thus allowing for a more precise understanding of the community^25^. Using a 4-dimensional culture system measuring bioluminescence of PER2∷LUC and *Bmal1-ELuc* enteroids, we explored the relationship between the host circadian clock and ASF metabolites. Using machine learning, we shortlisted metabolite candidates from the ASF metabolome^25^ to develop a targeted empirical approach. Our findings revealed that SCFAs are key discriminating features amongst the ASF metabolome involved in causing a circadian phase delay. Additionally, we provide evidence of HDAC inhibition as a mechanism for circadian rhythm entrainment.

Initial evidence for the role of microbial metabolites on host circadian rhythms was uncovered when phase delays were seen in both PER2∷LUC and *Bmal1-ELuc* abundance in enteroids exposed to ‘shifter’ ASF bacterial supernatants. Using machine learning, we identified SCFAs as specific metabolites contributing to these phase shifts. Individually testing of numerous microbial metabolites poses a time-consuming challenge to identify key metabolites. By utilizing a systems biology approach, we were able to circumvent this constraint to shortlist discriminating metabolite features.

Based on these findings, we compared the effects of SCFAs acetate, butyrate, formate, isovaleric acid and propionate against non-SCFA metabolites such as nicotinamide, choline and L-glutamine to validate this model. Nicotinamide, choline and L-glutamine are metabolites differentially utilized by the various ASF species and thus appropriate controls for SCFAs. The SCFAs acetate, butyrate, propionate and isovalerate caused pH-independent, dose-dependent phase delays whereas the SCFA formate and the remaining metabolites did not. With these findings we can confidently state that the key components of microbe-produced metabolites involved in regulating and entraining host circadian rhythms are SCFAs. Interestingly, the role of butyrate and acetate related to circadian rhythms has been described in previous literature with evidence of circadian phase alteration in mouse peripheral tissue due to oral administration of SCFAs^46^. However, the role of isovalerate has been limited to studies related to insomnia and depression^47^.

Core circadian genes are controlled through a variety of pathways and impact a diverse repertoire of processes, such as immune response, metabolism, and motility^48^. Several pathways may play a role in SCFA-induced clock entrainment. Pathways of interest involve SCFA-induced HDAC inhibition. SCFAs have been reported as potent HDACis with important effects on histone modifications^36,49,50^ including a recent study demonstrating that gut microbiota can influence HDAC activity via microbial-derived metabolites such as butyrate and valerate^49^. A recent study has also shown that the intestinal microbiota can mediate daily metabolic cycles epigenetically via the rhythmic induction of HDACs in epithelial cells of the small intestine^22^ whereas previous studies have suggested the importance of HDACs in regulating circadian rhythms of hepatic lipid metabolism^20^ and their roles in diet induced obesity and rhythmicity of the lipidome^51,52^. Therefore, we hypothesized that HDAC inhibition may play an important role in SCFA-induced circadian rhythm entrainment.

In support of this hypothesis we demonstrated that the HDACis TSA and SAHA mimic the effects seen by 1mM of butyrate on PER2∷LUC abundance. Additionally we showed PER2∷LUC phase delays with HDACis LP-411, HDAC1-2 inhibitor, MS275 and a botanic, synbiotic HDAC inhibitor, sulforaphane suggesting that entrainment of host circadian rhythms likely involves an HDAC inhibitor mechanism. Subsequent inhibition of HDAC inhibition using Mithramycin A reduced butyrate-induced PER2∷LUC phase delay confirming this hypothesis.

Further, by exposing human *Bmal1-luc* enteroids to either ASF356 supernatant or propionate we were able to confirm more modest phase delays in *Bmal1-luc* expression. Timing is a fundamental component of precision medicine. The presence of robust circadian rhythms in human enteroids will likely prove to be a valuable feature for future studies of the temporal dynamics of host-microbe interactions and for developing chronotherapeutic approaches targeted to the intestinal epithelium. Although beyond the scope of the current study, the contrasting nocturnal and diurnal activities of the murine and human intestinal epithelium may persist *ex vivo*.

With these findings we illuminate a novel mechanism by which the gut microbiota plays a role in maintaining and altering intestinal circadian rhythms. SCFAs and HDACis have been used for their anti-tumor and anti-inflammatory properties and for ameliorating conditions such as diversion colitis^53^ and inflammation in mouse models of DSS colitis^54,53,55,56^. Others have shown depletion of colonic SCFAs increases susceptibility to colitis^57^. HDACs repress transcription by deacetylating histones but also target nonhistone proteins,^22^ thereby affecting gene expression. Butyrate, for instance, increases differentiation of colonic T cells by modified histone acetylation of the *Foxp3* promoter, leading to protection against colitis^58,59^. Additionally, sulforaphane acts through a number of signaling pathways including NF-κB and Nrf2 (nuclear factor erythroid 2– related factor 2)^60,61^.

Our study sets the stage for potential translational applications. With the advent of chronotherapy^62^, potential therapeutic avenues to explore including: 1) using ‘shifting’ metabolites to prime the intestinal clock to decolonize invasive bacteria or prepare for chemotherapy, 2) treatment of jet lag and sleep disorders and acclimation of shift workers through microbial metabolite-mediated host clock entrainment, ^63–67^. Interestingly, isovaleric acid (a metabolite also found in valerian, a traditional sleep aid and anxiolytic ^68^) induced an increase in the amplitude of PER2∷LUC oscillations. Previous studies have shown circadian gene amplitude increases help protect against metabolic syndrome^69^. With further investigation, circadian rhythm modulation via the microbiome may play an important role in treating disease.

Enteroid models possess multiple gastrointestinal cell types and provide a unique model for understanding the relationship between the host and microbiome in relation to circadian rhythms. However, due to their complexity, certain limitations arise. We did not establish the mechanistic role for specific intestinal epithelial cell types; however, our previous studies have identified Paneth cells as key circadian pacemakers ^24^. In this study, we exposed the basolateral surface of 3D enteroids to metabolites. *In vivo*, these metabolites would be more concentrated at apical surfaces. In future studies, it will be equally important to investigate SCFA exposure in colonoids and enteroids. Colon epithelial cells have greater exposure to gut bacteria and utilize SCFAs as a fuel source^70^. Interestingly, we note that 3 of 4 ASF shifter species (ASF356, ASF492 and ASF500) consistently maintain low levels in the small intestine and show a sharp increase in abundance at the ileocecal valve^71^. Further, it has been shown that ASF519, which induced the strongest phase shifts in our experiments, is the most dominant ASF species in several mouse models^45^. Hence, host phase shifts may prove to be an intriguing mechanism by which certain bacterial species establish their niche within the gut. We observed changes in amplitude of clock genes, but did not measure the abundance of these genes by qRT-PCR to determine whether expression or overall abundance of these genes was affected. Lastly, we investigated contributions of individual ASF species on the clock, however more work is needed to determine how complex communities of microbes impact host circadian rhythms in aggregate.

Future studies will also require a deeper understanding of the importance of microbe-mediated host circadian rhythms in gastrointestinal pathophysiology. Butyrate is a well established therapy for diversion colitis and recent studies highlighted the role of gut commensal-produced propionate in mitigating colonization and invasion of *Salmonella*^72,73^. Given the impact of SCFAs on the host clock, further questions should focus on the extent to which these benefits require alteration of the host clock. Several gut immune cell populations are also under circadian control^1^. Intestinal epithelial organoids cocultured with immune cells would provide a more diverse model of circadian gut immunity. Further studies are also needed to uncover the evolutionary benefits of host clock alteration by gut microbes. Do SCFAs coordinate host metabolism or defense in coordination with host feeding? Does SCFA production allow SCFA producers to manipulate the host clock to favor their niche within the gut?

In conclusion, our study demonstrates gut microbe-generated metabolites entrain host circadian rhythms via an HDAC inhibition mechanism. Additionally, we provide new evidence and a systems based approach for linking individual bacterial taxa to specific microbial metabolites to and the host circadian clock.

## ACKNOWLEDGMENTS

We thank Gregory Medlock and Matthew Biggs from the Papin lab at UVA for providing ASF bacteria for generating bacterial supernatants, Glynis Kolling and Casandra Hoffman for advice and feedback, and Lubaina Ehsan for the graphical abstract. We also thank Drs. David Welsh at University of California, San Diego and Yoshihiro Nakajima at National Institute of Advanced Industrial Science and Technology for *Bmal1-Eluc* mice.

