## Supplementary material and figures for "Histone deacetylase inhibition by gut microbe-generated short chain fatty acids entrains intestinal epithelial circadian rhythms"

**Media Formulations**

Supplemented Brain–Heart Infusion medium (BHI) was prepared using BHI base (37 g/L, BD, Franklin Lakes, NJ, USA), supplemented with yeast extract (5 g/L), 0.2 mL of vitamin K1 solution (0.5% vitamin K1 dissolved in 99.5% ethanol), 0.5 mL/L of hemin solution (0.5 g/L dissolved in 1% NaOH, 99% deionized water), L-cysteine (0.5 g/L) and 5% each of newborn calf serum, horse serum, and sheep serum. Vitamin K1, hemin, and all sera were added after autoclaving the medium. For preparation of agar plates, agar was supplemented at 12g/L.

Modified Lennox Broth medium (mLB) was prepared using 30 g/L LB base in powder form (Sigma, St Louis, MO, USA), 0.376g/L L-cysteine (Sigma), 39mL of a mineral salts solution (containing 6 g/L KH_2_PO_4_, 6 g/L (NH_4_)_2_SO_4_, 12g/L NaCl, 2.5g/L MgSO_4_,7H2O, 1.6 g/L CaCl_2_,2H_2_O, dissolved in DI water), 15 mL/L of hemin solution (0.5 g/L dissolved in 1% NaOH, 99% DI water), 0.3 mL of vitamin K1 solution (50 mg/mL vitamin K1 dissolved in 200 proof ethanol), 15 mL/L of lactose solution (5g/L lactose dissolved in DI water) and 15 mL/L of tween 20 solution (1 g/L tween 20 dissolved in DI water).

L-WRN conditioned medium was prepared as previously described using L-WRN (ATCC® CRL-3276™) cells that secrete growth factors essential for gastrointestinal stem cell propagation ^25^. Basic minigut medium was prepared by combining 95mL of DMEM/F12, 1mL L-Glutamine, 1mL Pen/Strep (1:100), 1mL HEPES (1M), and 2mL B27 supplement (1:50). Murine Enteroid Growth Medium (mEGM) was prepared by mixing 30% of L-WRN conditioned medium with 70% of basic minigut medium. Human *Bmal1-luciferase (Bmal1-luc)* enteroids were grown in Stemcell IntestiCult™ Organoid Growth Medium (Human) (OGM) (CAT#06010).

**Human *Bmal1-luc* enteroid generation**

Plasmid DNA was packaged into lentiviral vectors by the Viral Vector Core at Cincinnati Children’s Hospital Medical Center (CCHMC). The protocol for transducing human intestinal enteroids (HIEs) was made by merging a HIE digestion ^31^ and mouse enteroid transduction protocol ^32^. One high density-well of enteroids was collected in a 1.5mL Eppendorf and pelleted three times, serially, to fully remove Matrigel. HIEs were digested to single cells via treatment with 400μL 0.5mM EDTA/0.05% Trypsin solution (Gibco) at 37°C for 5 minutes. Trypsin was deactivated with 900μL Advanced DMEM F12 (Gibco) supplemented with 10% Fetal Bovine Serum (FBS) (Gibco). Cell clumps were further digested with repeated pipetting using a p1000 pipette. Single cells were pelleted and resuspended in viral media containing: 50μL Lentiviral suspension (titer ≈10^6-7^) (CCHMC), 8μg/mL Polybrene (Millipore-Sigma), 10μM CHIR99021 (Cayman), 10μM ROCK inhibitor (Millipore-Sigma) and Intesticult Component A/B (1:1 mix) to bring the volume to 400μL. Viral media suspensions were added to one well of a 24-well cell culture dish and spinoculated for 1-hour x600g at 32°C. The plate was then placed in a 37°C, 5% CO2 cell culture incubator for 6-hours. Following incubation, cell suspensions were added to a 1.5mL Eppendorf, pelleted, suspended in 30μL of Matrigel, and plated as three 10μL domes/well in a new 24-well plate. 350μL of Intesticult Component A/B (1:1) media supplemented with 10μM ROCK inhibitor and 10μM CHIR99021 was added to the culture after Matrigel solidification. After 24-hours, media was replaced with 350μL Intesticult Component A/B (1:1) supplemented with 10μM ROCK inhibitor. On day two post-infection, media was transitioned to Intesticult Component A/B (1:1) supplemented with 2μg/mL Puromycin. Puromycin selection was maintained for 2-weeks before starting experiments.

**Supplementary Figures**

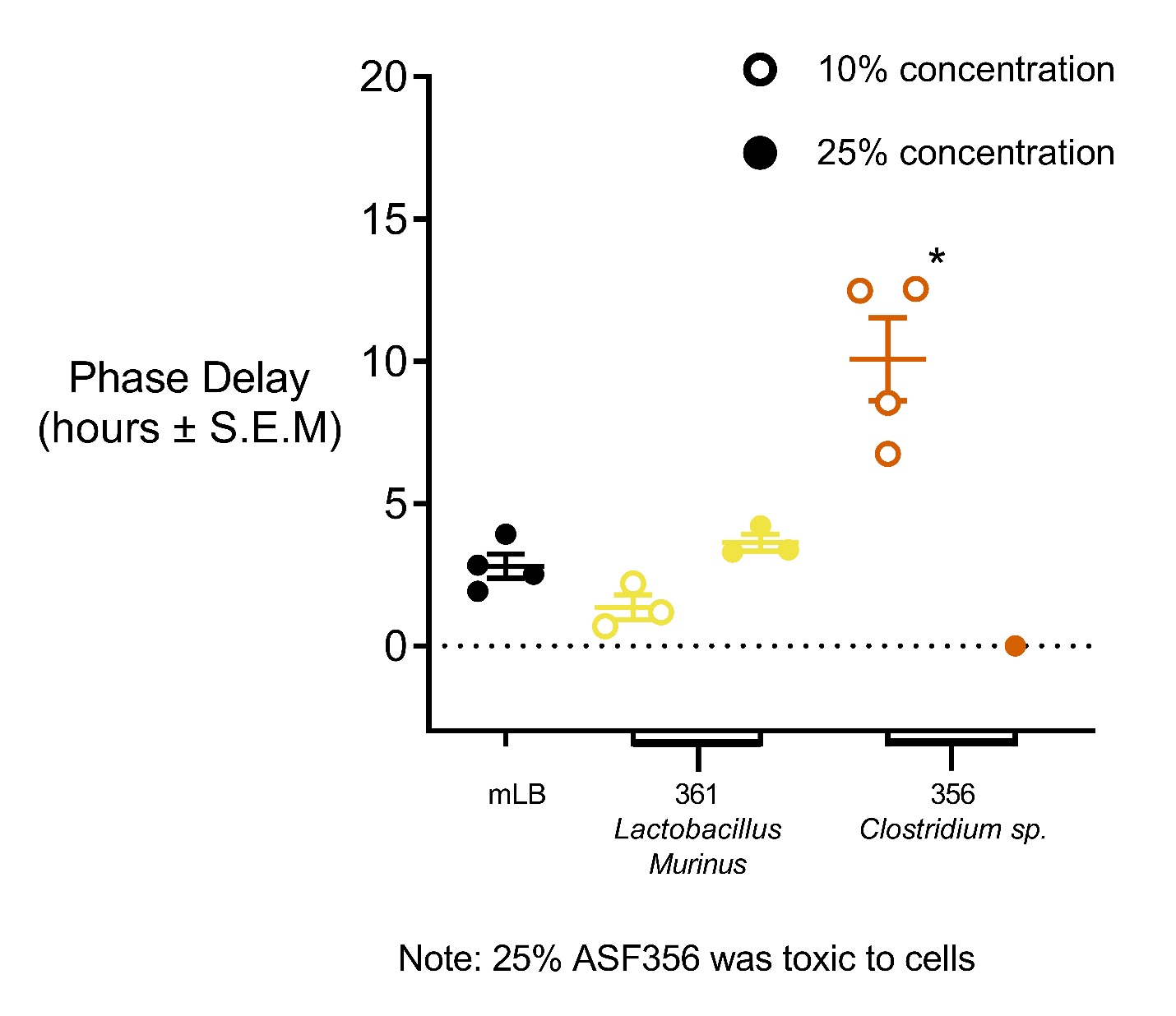

Supplementary Figure 1

Median phase delay (represented by middle horizontal line of samples) due to mLB or ASF bacterial supernatant calculated as a forward shift in *Bmal1-Eluc* oscillation compared to an untreated control. ASF ‘shifters’ classified as ASF taxa causing statistically significant phase delays of PER2::LUC oscillation as compared to a 25% mLB control (*P<0.05, Mann Whitney *U* test). 25% ASF356 was toxic to enteroids.

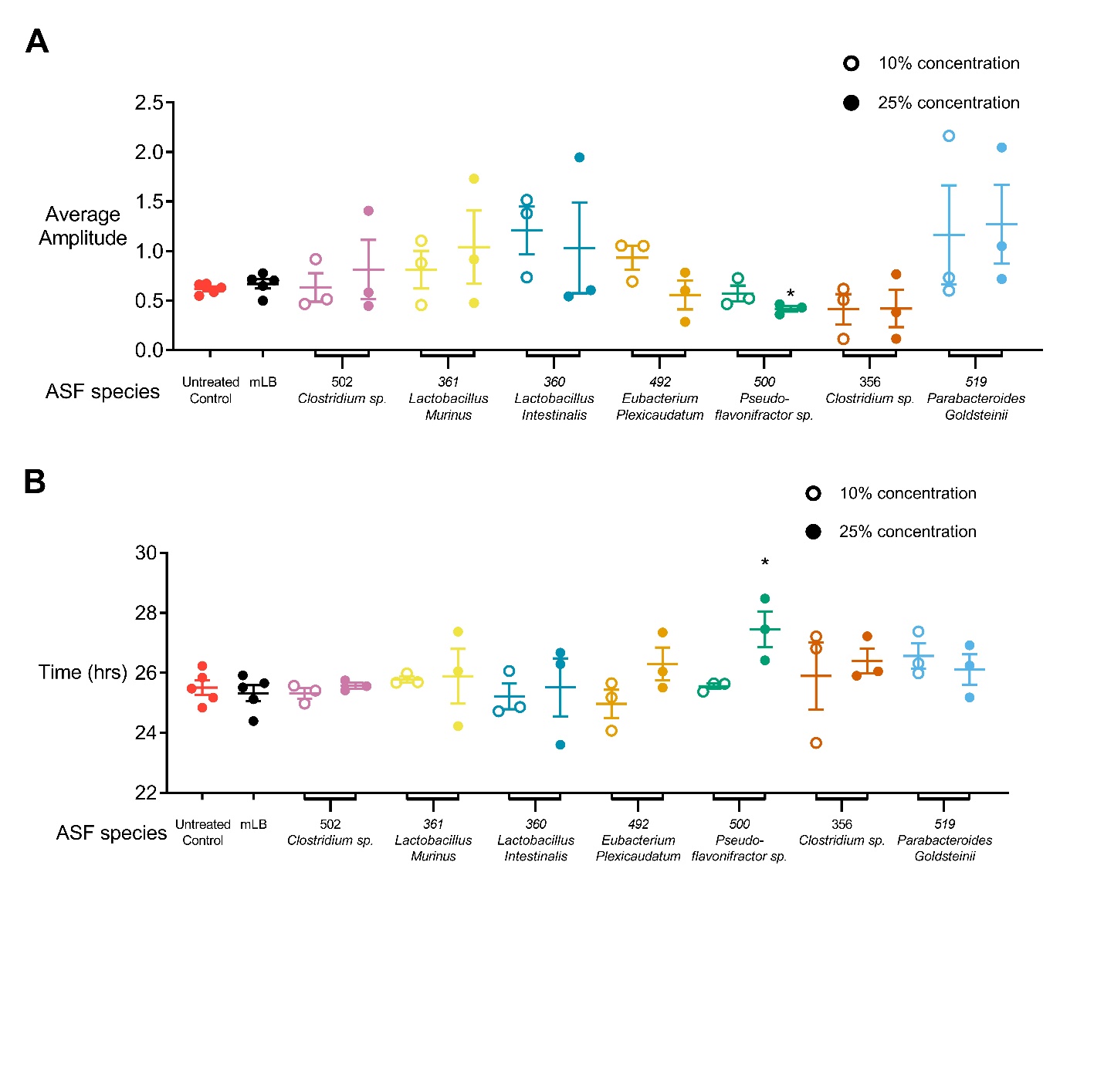

Supplementary Figure 2

A) Median amplitude (represented by middle horizontal line of samples) of PER2::LUC abundance in presence of either mLB or ASF bacterial supernatant calculated as a normalized maximum luminescence of PER2::LUC abundance. PER2::LUC oscillation peaks in presence of ASF bacterial supernatant were compared to 25% mLB control. (*P<0.05, Mann Whitney U test). 25% of ASF500 led to a dampening of amplitude of PER2::LUC abundance B) Median period (represented by middle horizontal line of samples) of PER2::LUC oscillations in presence of either mLB or ASF bacterial supernatant calculated as time for a complete PER2::LUC oscillation to complete. PER2::LUC oscillation periods in presence of ASF bacterial supernatant were compared to 25% mLB control. (*P<0.05, Mann Whitney U test).

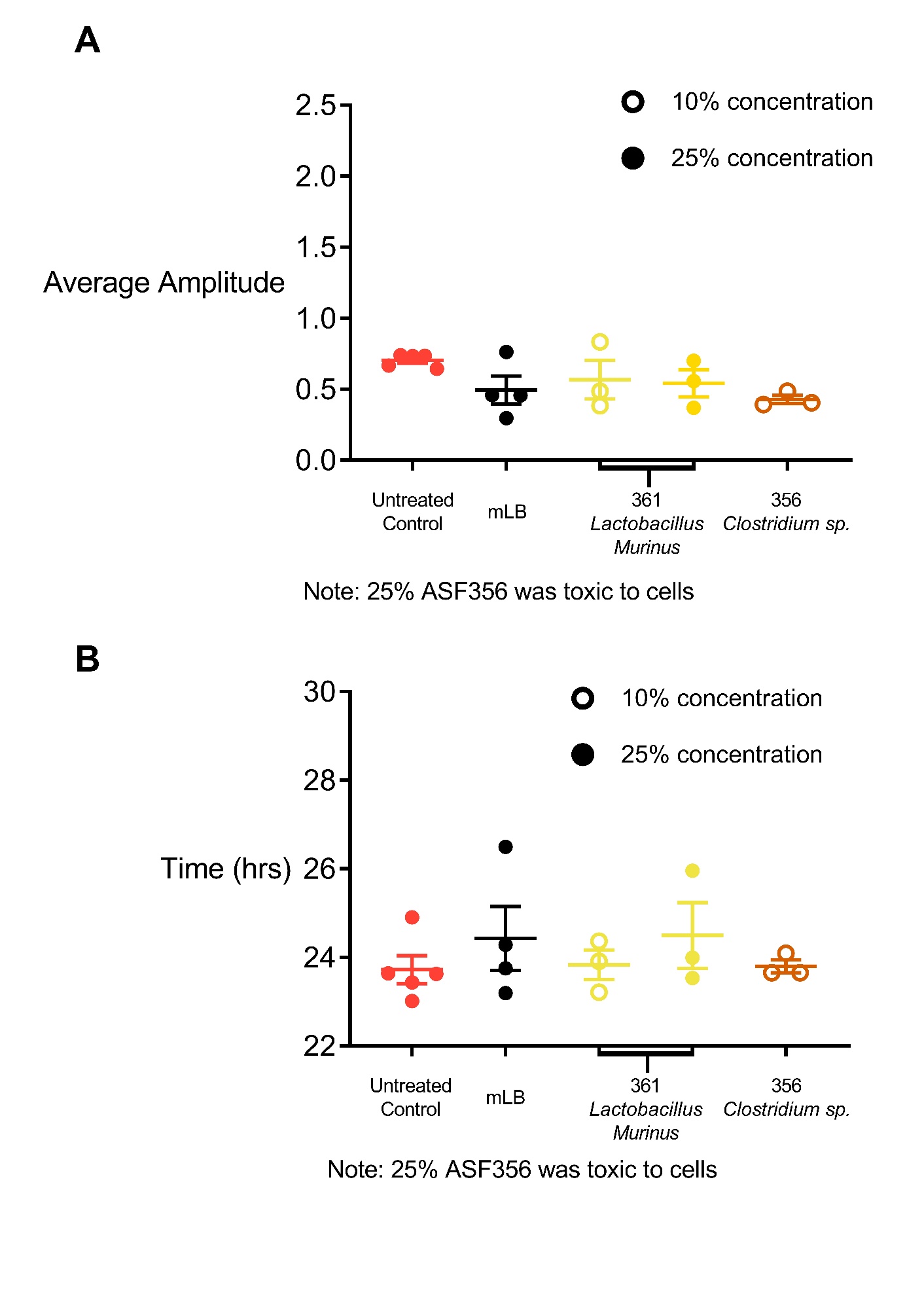

Supplementary Figure 3

A) Median amplitude (represented by middle horizontal line of samples) of *Bmal1-Eluc* abundance in presence of either mLB or ASF bacterial supernatant calculated as a normalized maximum luminescence of *Bmal1-Eluc* abundance. *Bmal1-Eluc* oscillation peaks in presence of ASF bacterial supernatant were compared to 25% mLB control. (Mann Whitney U test). B) Median period (represented by middle horizontal line of samples) of *Bmal1-Eluc* oscillations in presence of either mLB or ASF bacterial supernatant calculated as time for a complete *Bmal1-Eluc* oscillation to complete. *Bmal1-Eluc* oscillation periods in presence of ASF bacterial supernatant were compared to 25% mLB control. (Mann Whitney U test).

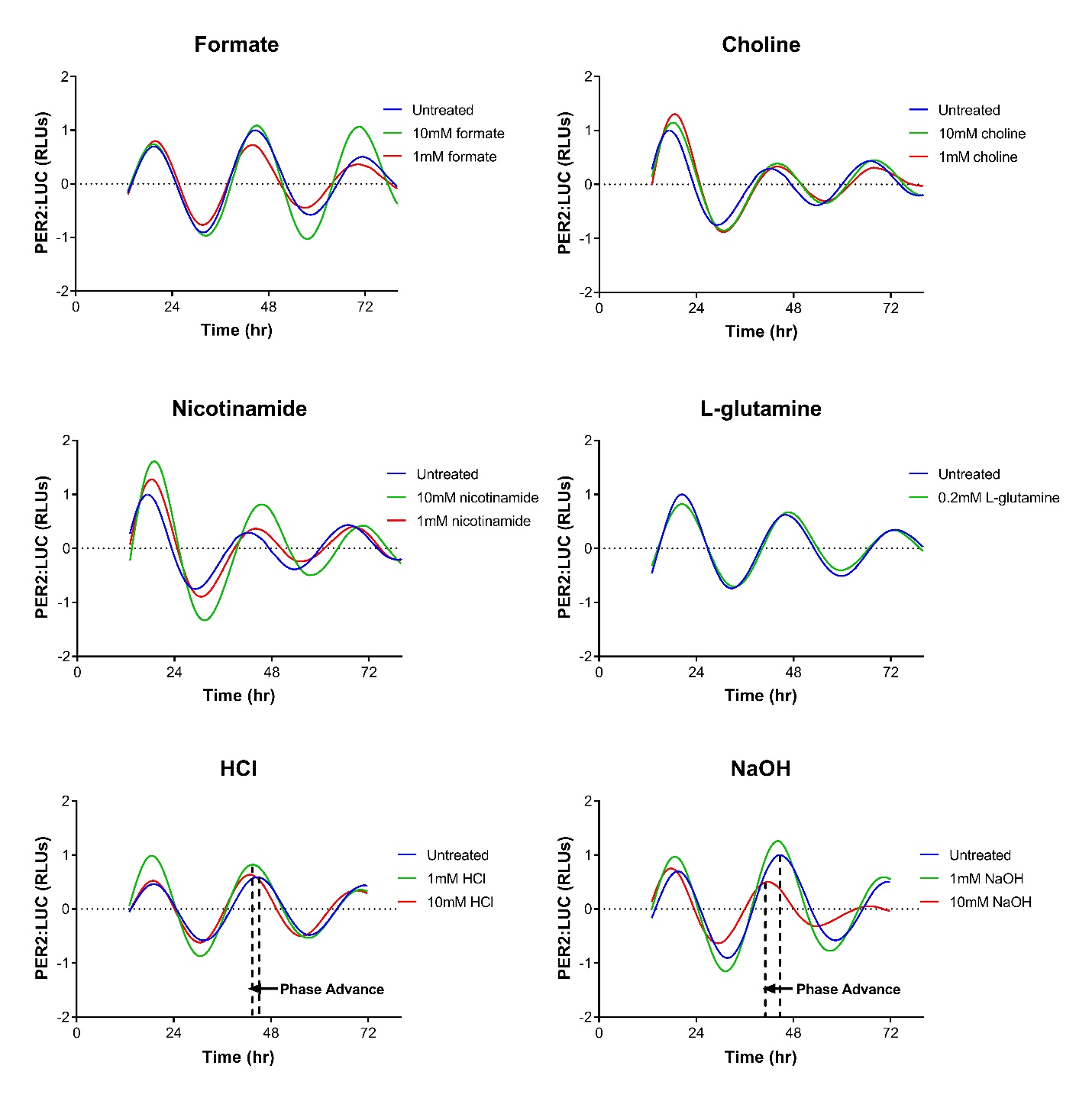

Supplementary Figure 4

Representative PER2::LUC enteroids displaying PER2::LUC oscillations over 72 hours. Formate, choline, nicotinamide and L-glutamine do not show evidence of phase delay while hydrochloric acid (HCl) and sodium hydroxide (NaOH) demonstrate phase advances, represented as a backward shift in the PER2::LUC waveform (P<0.05 by Mann-Whitney *U* test, n=3 samples per metabolite).

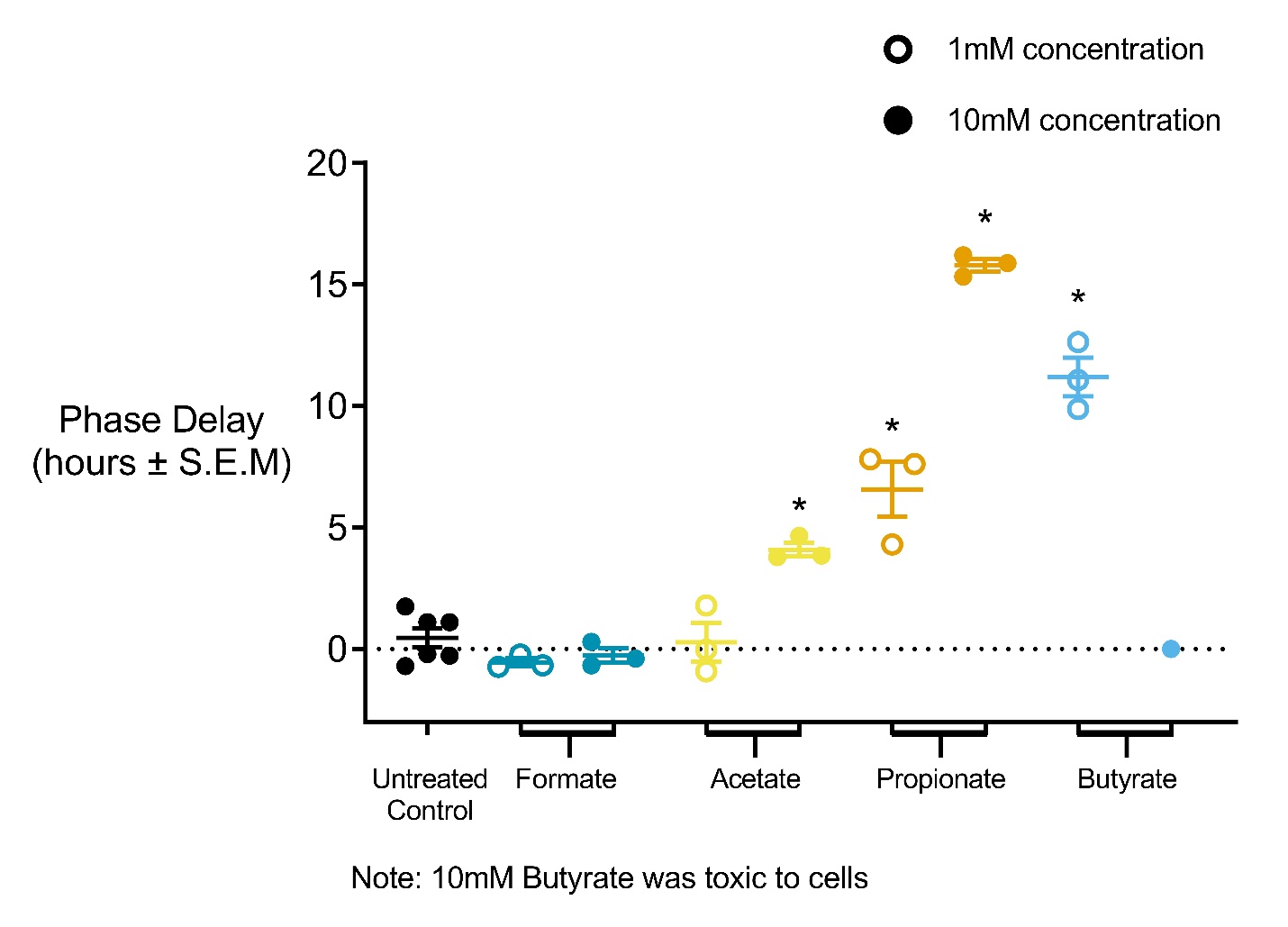

Supplementary Figure 5

Median phase delay (represented by middle horizontal line of samples) due to metabolites calculated as a forward shift in *Bmal1-Eluc* oscillation compared to an untreated control (*P<0.05, Mann Whitney *U* test). 10mM acetate, 1mM propionate, 10mM propionate and 1mM butyrate caused significant phase delays of *Bmal1-Eluc* abundance. 10mM butyrate was toxic to enteroids.

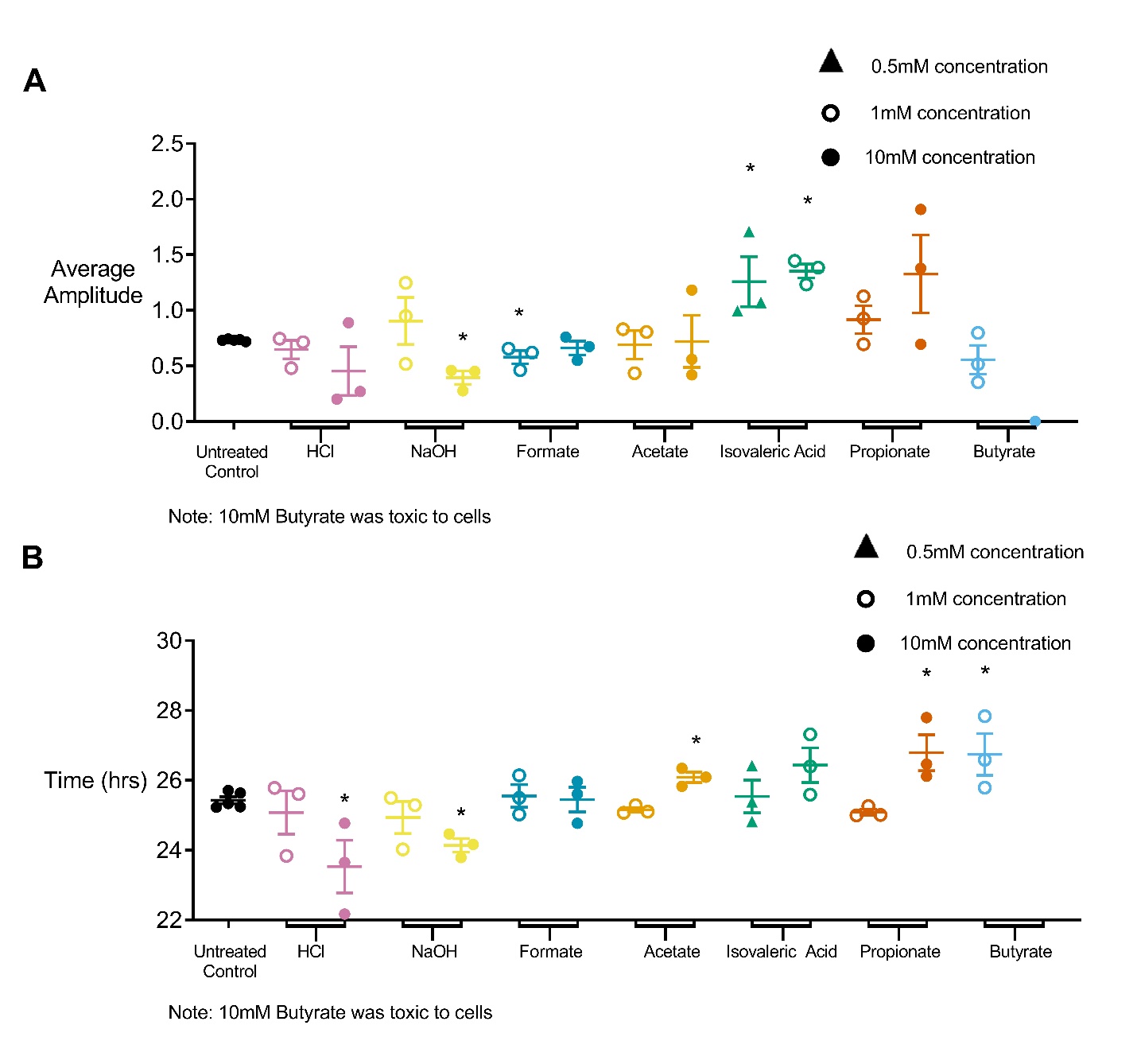

Supplementary Figure 6

A) Median amplitude (represented by middle horizontal line of samples) of PER2::LUC abundance in presence of metabolites calculated as a normalized maximum luminescence of PER2::LUC abundance. PER2::LUC amplitudes in presence of metabolites were compared to an untreated control. (*P<0.05, Mann Whitney U test). 10mM sodium hydroxide (NaOH) and 1mM formate led to a dampening of amplitude of PER2::LUC abundance while both 0.5mM and 1mM isovaleric acid showed significant increases in amplitude B) Median period (represented by middle horizontal line of samples) of PER2::LUC oscillations in presence of metabolites calculated as time for a complete PER2::LUC oscillation to complete. PER2::LUC oscillation periods in presence of metabolites were compared to an untreated control. (*P<0.05, Mann Whitney U test). 10mM hydrochloric acid (HCl) and 10mM NaOH decreased PER2::LUC period while 10mM acetate, 10mM propionate and 1mM butyrate increased PER2::LUC period.

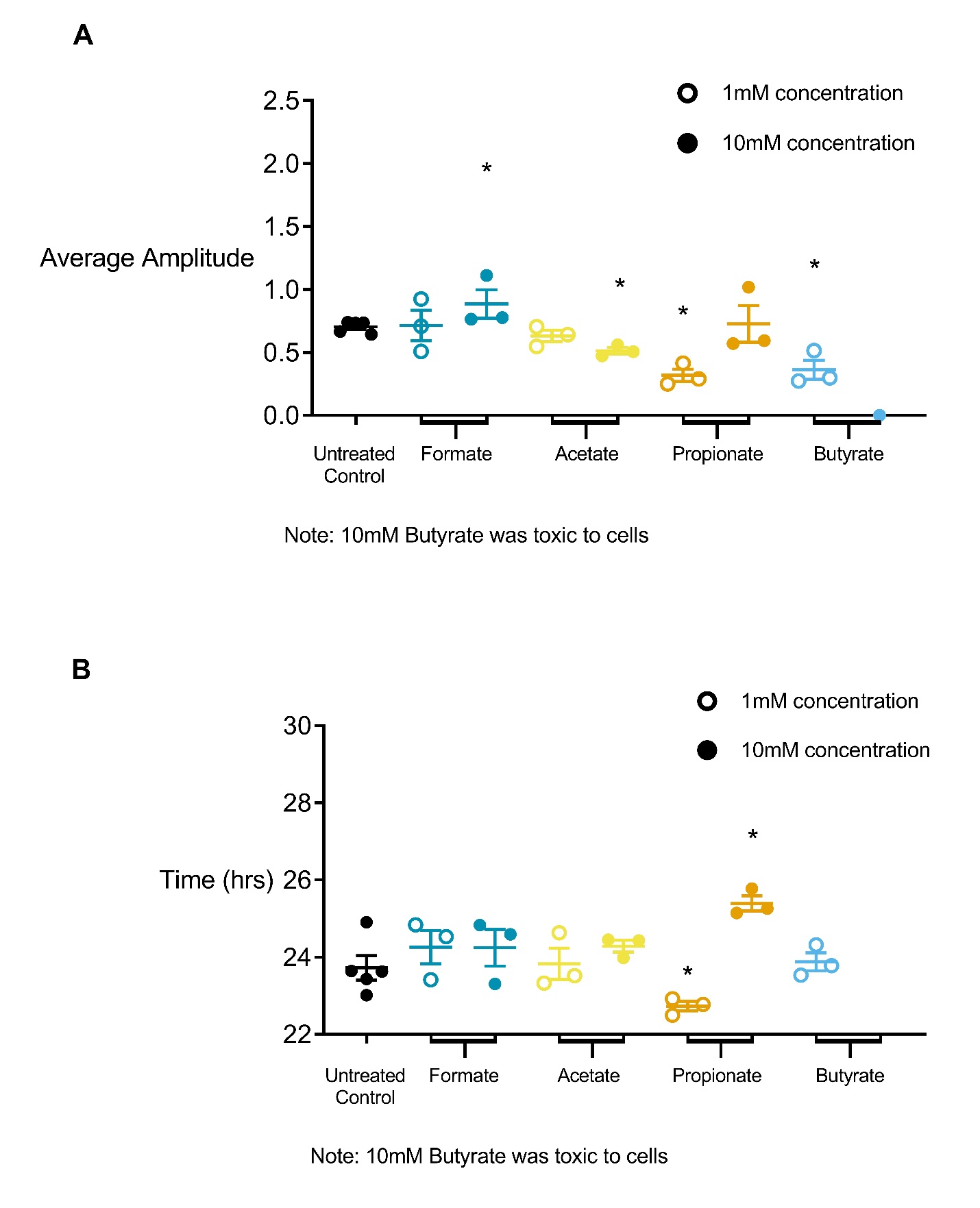

Supplementary Figure 7

A) Median amplitude (represented by middle horizontal line of samples) of *Bmal1-Eluc* abundance in presence of metabolites calculated as a normalized maximum luminescence of *Bmal1-Eluc* abundance. *Bmal1-Eluc* amplitudes in presence of metabolites were compared to an untreated control. (*P<0.05, Mann Whitney U test). 10mM acetate, 1mM propionate and 1mM butyrate led to a dampening of amplitude of PER2::LUC abundance while both 10mM formate showed a significant increase in amplitude B) Median period (represented by middle horizontal line of samples) of *Bmal1-Eluc* oscillations in presence of metabolites calculated as time for a complete *Bmal1-Eluc* oscillation. *Bmal1-Eluc* oscillation periods in presence of metabolites were compared to an untreated control. (*P<0.05, Mann Whitney U test). 1mM propionate decreased *Bmal1-Eluc* period while 10mM propionate increased *Bmal1-Eluc* period.

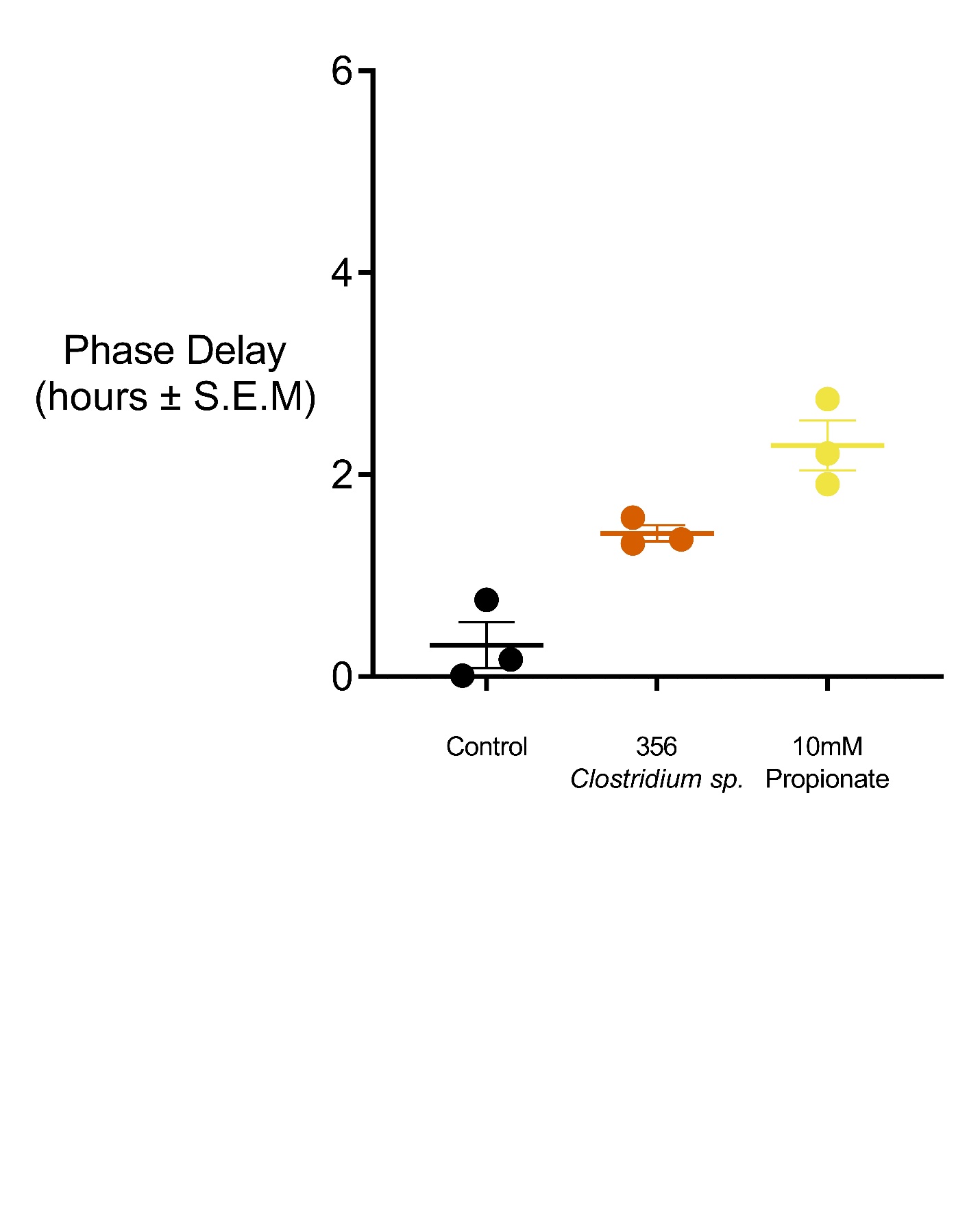

Supplementary Figure 8

Median phase delay (represented by middle horizontal line of samples) due to ASF bacterial supernatant calculated as a forward shift in human *Bmal1-luc* oscillation compared to an untreated control. ASF bacteria show trends in phase delays of *Bmal1-luc* abundance when compared with an untreated control.
